# A critical role of epithelial MHCII in initiation of autoimmune tumorigenesis and sustaining premalignancy growth in the stomach

**DOI:** 10.64898/2026.04.01.715843

**Authors:** Clarisel C. Lozano, Emily N. Vazquez, Alexander Kolev, Amanda M. Honan, Wael El-Rifai, Alexander Zaika, Zhibin Chen

## Abstract

Autoimmunity is emerging as a new etiology for early-onset gastric cancer (GC). However, it remains unclear what molecular pathways drive the initiation and progression of autoimmune tumorigenesis. Given that Major Histocompatibility Complex Class II (MHCII) is the strongest genetic risk factor for many autoimmune diseases, we hypothesized that MHCII-mediated autoantigen presentation drives tumorigenic differentiation of epithelial cells. Here we show that epithelial MHCII, rather than MHCII from immune cells, plays an essential role in the initiation of autoimmunity-driven tumorigenic differentiation of gastric epithelial cells, which was characterized by increased expression of cancer-associated markers with immune-evasive and stem-like features that potentiate premalignant progression. In addition, we show that early gastric premalignancy is reversible upon the removal of epithelial MHCII. This study reveals that epithelial MHCII antigen presentation is essential in the early stages of autoimmune-driven gastric tumorigenesis and highlight epithelial MHCII as a potential biomarker or therapeutic target in early interventions of autoimmunity-driven cancer development.

## Introduction

Gastric cancer (GC) is the fifth leading cause of cancer-related mortality worldwide, with almost 1 million new cases reported annually, and more than 650,000 deaths globally^1^. The incidence of early-onset GC (diagnosis in people younger than 50 years) is rising, with a disproportionate increase observed in females relative to males. *Helicobacter pylori* infection, as well as environmental and host-intrinsic risk factors, such as genetics and nutrition, are strong risk factors for gastric cancer. Across these diverse etiologies, chronic mucosal inflammation emerges as a shared driver of premalignant transformation, implicating immune dysregulation as a central mechanism in gastric carcinogenesis.

Gastric adenocarcinomas (GA) arise from gastric epithelial cells and account for more than 90% of all gastric cancer cases^2^. The Correa cascade has been established as a model outlining the stepwise progression from normal gastric mucosa to cancer through the following histopathological stages: chronic gastritis, gastric atrophy, intestinal metaplasia (IM), dysplasia, and ultimately invasive gastric cancer^1,3^. Spasmolytic polypeptide-expressing metaplasia (SPEM) has been identified as a possible precursor lesion of IM in human GA, though the precise relationship between SPEM and IM remains an area of active investigation^4^. Inflammatory signals have been shown to be required for SPEM or IM progression to dysplasia, in both human disease and mouse models^5^. SPEM development reflects phenotypic reprogramming of the gastric epithelium in response to chronic inflammation, which may originate from bacterial infections such as *H. pylori*. These inflammatory signals may also arise from autoimmune responses, for example, in autoimmune gastritis (AIG), in which T cell-mediated destruction of parietal cells drives chronic inflammation, representing a clinical context where SPEM development occurs independently of bacterial infection ^6,7^.

CTLA4, an immune checkpoint receptor that maintains peripheral T cell tolerance, has emerged as a risk factor for gastric cancer. In humans, *CTLA4* haploinsufficiency led to GC in 12.5% of reported cases (3 of 24 patients), with 2 of those 3 patients presenting with adenocarcinoma associated with atrophic gastritis and intestinal metaplasia ^8^. The consistent association with premalignant histopathology demonstrates a mechanistic link between *CTLA4* insufficiency and epithelial transformation.

To model CTLA4 haploinsufficiency, our lab developed a transgenic CTLA4 shRNA (CTLA4KD) mouse model that reduces expression of CTLA4 by ∼60%. Mice carrying the CTLA4KD recapitulate the Correa cascade of gastric cancer, developing autoimmune gastritis and SPEM by 5-6 weeks of age, and progressing to gastric adenocarcinoma after 12 months of age^9^. To our knowledge, this represents the only genetically defined murine model that faithfully recapitulates the progression from autoimmune-driven gastritis to invasive adenocarcinoma, providing a system to interrogate the inflammatory signals that initiate and sustain premalignant transformation. Previous work from our laboratory established that CD4 T cells are necessary and sufficient to drive SPEM development in CTLA4KD mice, yet the molecular mechanism by which autoreactive CD4 T cells instruct the gastric epithelium to undergo premalignant reprogramming remain unknown.

As the global burden of autoimmune disease increases, the genetic determinants of autoimmune susceptibility have become an area of intense investigation. For several autoimmune conditions, including type 1 diabetes (T1D), multiple sclerosis (MS), rheumatoid arthritis (RA), and coeliac disease, the major histocompatibility complex class II (MHCII; human leukocyte antigen, HLA class II) carries the strongest known genetic determinants of disease risk ^10,11^. Therefore, we adapted the CTLA4KD model to investigate the role of MHCII in autoimmune-driven gastric carcinogenesis. Here, we demonstrate that epithelial MHCII, rather than MHCII expression on professional antigen-presenting cells, plays a critical role in initiating autoimmunity-driven premalignant transformation. Epithelial MHCII expression was associated with expansion of gastric epithelial populations bearing cancer progenitor markers and immune-evasive and stem-like features that potentiate premalignant progression. Strikingly, early gastric SPEM was reversible upon conditional removal of MHCII, demonstrating the dependency of the premalignant program on this epithelial signal. Collectively, this study identified a novel role for epithelial MHCII in driving premalignant differentiation in the gastric epithelium and establishes epithelial antigen presentation as a previously unrecognized checkpoint in the transition from autoimmune inflammation to gastric premalignancy.

## Results

### Epithelial MHCII promotes initiation of autoimmune gastric tumorigenesis

To determine whether epithelial MHCII expression changes during the progression from premalignancy to adenocarcinoma, we analyzed gastric epithelial cells from CTLA4KD mice at premalignant and adenocarcinoma stages alongside age-matched CB6F1 controls using 40-color spectral flow cytometry. CTLA4KD mice on the Balb/c x C57BL/6 F1 (CB6F1) background develop autoimmune gastritis and severe metaplastic lesions by 5-6 weeks of age, and spontaneously transition to invasive gastric adenocarcinoma after 12 months of age ^9^. We analyzed gastric epithelial cells (PDPN^-^CD31^-^EPCAM^+^; gated to exclude stromal and endothelial cells) from CTLA4KD mice at the premalignant (young) and adenocarcinoma (old) stages, as well as age-matched CB6F1 wildtype (WT) controls (Fig. 1A). The proportion of MHCII-expressing gastric epithelial cells was significantly elevated in the adenocarcinoma-stage CTLA4KD mice relative to both premalignant and WT controls. We next examined the co-expression of MHCII and PDL1 on gastric epithelial cells, given the well-established role of PD-L1 on tumor cells in suppressing anti-tumor immunity and promoting cancer progression^12,13^. CTLA4KD mice at both premalignant and adenocarcinoma stages exhibited elevated co-expression of MHCII and PD-L1 on gastric epithelial cells compared to WT controls (Fig. 1B). This data suggests that epithelial MHCII expression is progressively upregulated during autoimmune-driven premalignant transformation and adenocarcinoma development.

**Figure 1.**
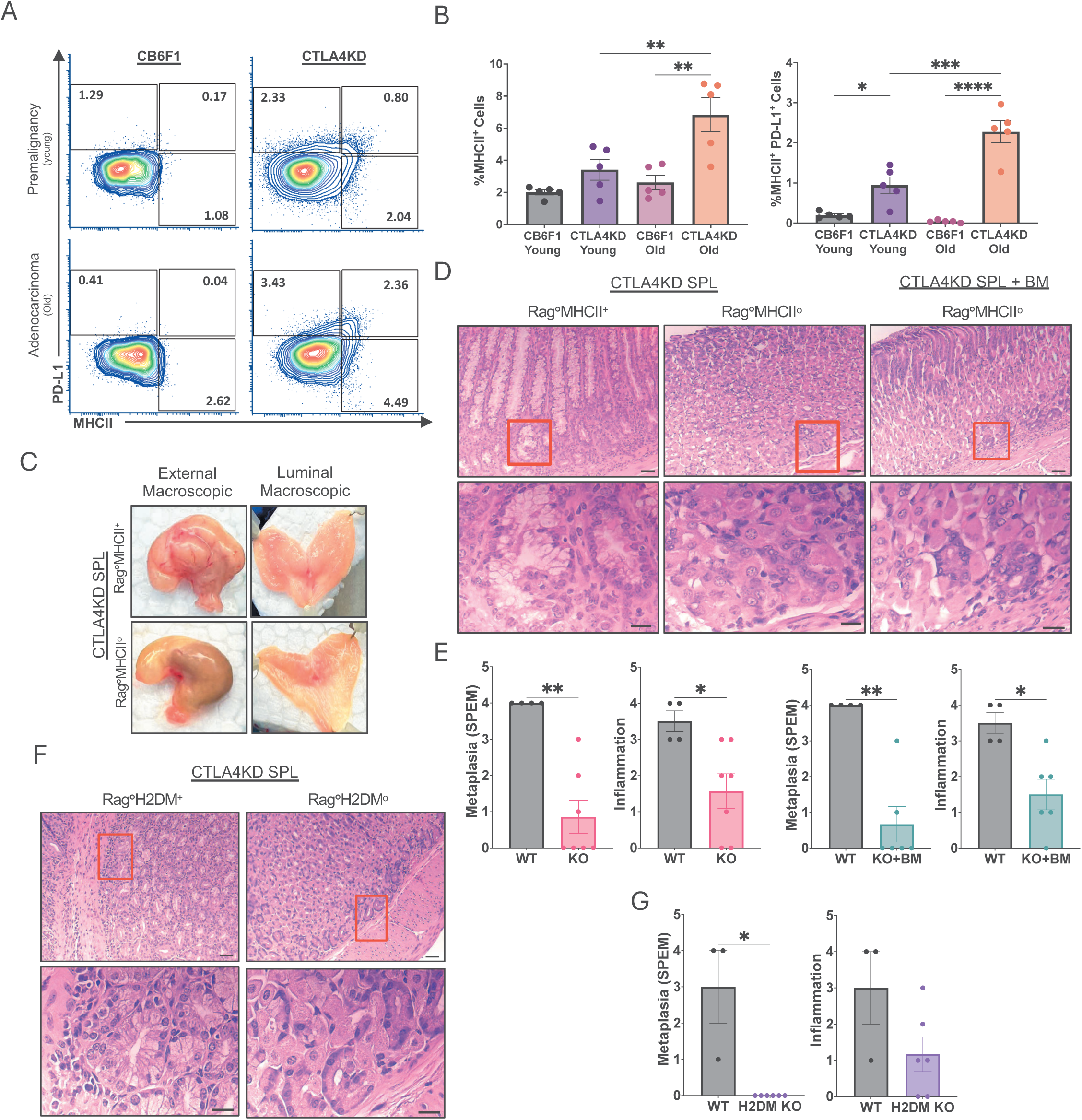
Requirement of MHCII for initiation of autoimmune driven gastric tumorigenesis. **(A)** Representative flow cytometry plots of PD-L1 and MHCII expression on gastric epithelial cells isolated from CTLA4 knockdown (CTLA4KD) mice at the premalignant (young) and adenocarcinoma (old) stages, and from age-matched CB6F1 controls. **(B)** Summary percentage of MHCII^+^ and MHCII^+^PD-L1^+^ gastric epithelial cells in CTLA4KD and CB6F1 age-matched cohorts. **(C)** Representative macroscopic photographs of the external gastric surface and luminal side from indicated experimental cohorts. **(D)** Representative hematoxylin and eosin (H&E)-stained gastric tissue sections at low (4X) and high (40X) magnification from Rag-deficient MHCII-knockout (Rag°MHCII°) and Rag-deficient MHCII-wildtype (Rag°MHCII°) mice. Hosts received adoptive transfer of CTLA4KD donor splenocytes, with or without co-transfer of Rag-deficient bone marrow (BM), at 3-10 days of age. Gastric tissue was analyzed 3 months post-transfer. **(E)** Histopathology scores for gastric inflammation and spasmolytic polypeptide-expressing metaplasia (SPEM) from cohorts described in (D). **(F)** Representative H&E-stained gastric sections at 4X and 40X magnification from Rag-deficient H2DM-knockout (Rag°H2DM°) and Rag-deficient H2DM-wildtype (Rag0H2DM^+^) mice following adoptive transfer of CTLA4KD donor splenocytes at 3-10 days of age, analyzed at 3 months post-transfer. **(G)** Histopathology scores for gastric inflammation and SPEM from cohorts described in (F). Scale bar, 4X:100 µm, and 40X: 50 µm. Each data point represents an individual mouse; (mean ± SEM; Mann-Whitney test), \**p* < 0.05, \*\**p* < 0.01, \*\*\**p* < 0.001.

To functionally test the requirement for MHCII in disease initiation, we utilized a Rag knockout MHCII KO (Rag°MHCII°) adoptive transfer model. Donor CTLA4KD splenocytes were adoptively transferred into neonatal Rag°MHCII^+^ (MHCII WT) or Rag°MHCII° (MHCII KO) recipients at 3-5 days of age. Three months post-transfer, tissues were harvested for analysis. Gross examination revealed marked differences between MHCII WT and KO groups. Stomachs from MHCII WT recipients displayed a wrinkled external surface and enlarged rugae on the mucosal surface, consistent with the gastric enlargement and mucosal hypertrophy previously described in CTLA4KD mice with SPEM (Fig. 1C). Histopathological analysis confirmed extensive SPEM-associated premalignant changes in MHCII WT recipients, whereas MHCII KO recipients showed absent or markedly reduced premalignant pathology (Fig. 1D, E). These findings establish that MHCII expression is required for SPEM development. However, because Rag°MHCII° hosts lack MHCII in both hematopoietic and non-hematopoietic compartments, this experiment alone cannot resolve whether the absence of MHCII on bone-marrow-derived APCs or epithelial cells is the primary determinant of premalignant gastric transformation.

To dissect the relative contributions of hematopoietic versus non-hematopoietic MHCII, Rag-deficient bone marrow (BM) from MHCII-wildtype donors was co-transferred with CTLA4KD splenocytes into neonatal MHCII KO recipients. This experimental design restores MHCII expression to BM-derived immune cells, including professional APCs such as dendritic cells and macrophages, while the non-hematopoietic compartment remains MHCII-deficient. Neonatal transfer allowed bone marrow reconstitution without the myeloablative irradiation required for adult recipients. MHCII KO recipients reconstituted with MHCII^+^ BM failed to develop characteristic SPEM pathology (Fig. 1D). Blinded histological scoring confirmed that both gastric inflammatory infiltration and SPEM severity were significantly reduced in MHCII KO and MHCII KO+BM groups relative to MHCII WT controls (Fig. 1E). Together, these data suggest that MHCII expressed by BM-derived immune cells is insufficient to initiate autoimmune-driven SPEM, implicating the non-hematopoietic compartment as the critical source of disease-relevant MHCII. We note that because the transferred CTLA4KD splenocyte pool may contain residual MHCII-expressing hematopoietic cells, a formal contribution from this source cannot be entirely excluded.

### Functional antigen presentation by epithelial MHCII is required for autoimmune-driven gastric tumorigenesis

The pathways for MHC trafficking and antigen loading have been well characterized, and studies have shown that MHCII molecules can exist in an empty conformation on the cell surface. In humans, HLA-DP alleles carrying the 84Gly beta chain polymorphism fail to bind CLIP and can assemble as empty molecules capable of binding exogenously derived peptides at the cell surface^14^, and in murine models, empty MHCII molecules have been identified on the surface of spleen- and BM-derived dendritic cells^15^. The existence of surface MHCII that is not engaged in classical peptide presentation raises the question of whether MHCII surface expression alone, rather than functional antigen loading, could be sufficient to drive the SPEM pathology response in our model.

To test this hypothesis, we generated a Rag°H2DM^+^ or Rag°H2DM° mouse model. H2-DM heterodimers catalyze the removal of the invariant chain-derived CLIP peptide from MHCII and facilitate the loading of high-affinity antigenic peptides^16,17^. In the absence of H2-DM, MHCII molecules are predominantly occupied by CLIP and are unable to present antigenic peptides to T cells, providing a means to uncouple MHCII surface expression from functional antigen presentation^18^. We adoptively transferred CTLA4KD splenocytes into Rag°H2DM^+^ or Rag°H2DM° neonates. Rag°H2DM° mice failed to develop characteristic SPEM pathology (Fig. 1F). Notably, gastric inflammatory infiltration was not significantly different between Rag°H2DM^+^ or Rag°H2DM° recipients, indicating that immune cell recruitment to the gastric mucosa was preserved in the absence of functional antigen presentation, but was insufficient to drive SPEM pathology (Fig. 1F, G). Collectively, these results demonstrate that functional antigen presentation by epithelial MHCII, rather than hematopoietic MHCII or MHCII surface expression alone, is required to drive premalignant pathology.

### Epithelial MHCII expression drives expansion of cancer progenitor and immune-evasive epithelial populations

To characterize the non-hematopoietic gastric compartment, we performed unbiased SPADE clustering of 40-marker spectral flow cytometry data from Rag°MHCII^+^, Rag°MHCII°, and Rag°MHCII° + BM hosts. Two EpCAM^+^ epithelial clusters were identified, an MHCII-low cluster (EPCAM1) present across all groups, and an MHCII-high (EPCAM2) cluster that was selectively enriched in Rag°MHCII^+^ hosts and virtually absent in Rag°MHCII° and Rag°MHCII° + BM hosts, confirming epithelial compartment-specific MHCII expression at single-cell resolution (Fig. 2A). These findings suggest that the EpCAM2 cluster represents an MHCII-expressing epithelial subset with putative antigen-presenting capacity that is selectively expanded in the context of autoimmune gastric inflammation.

**Figure 2.**
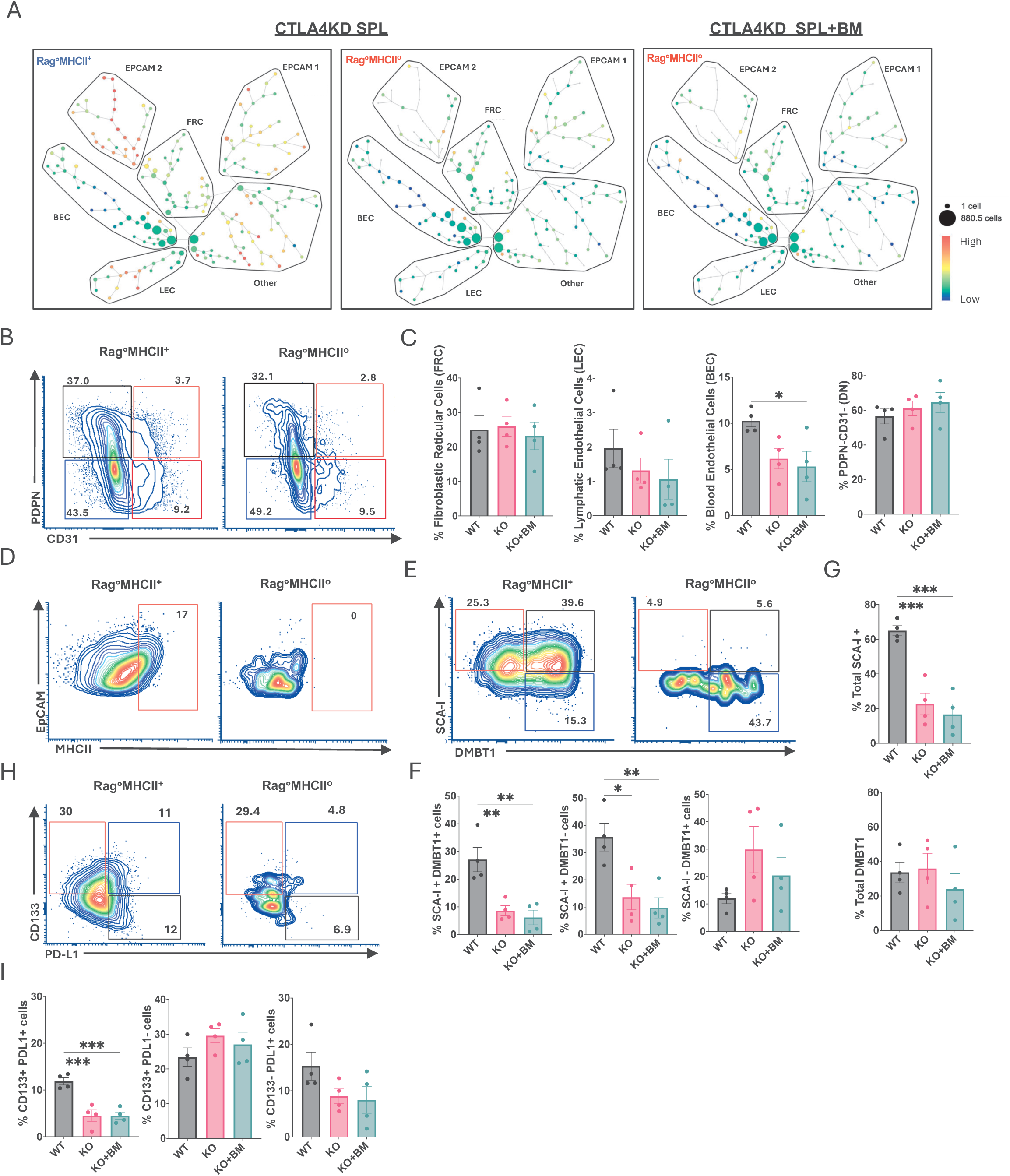
MHCII-dependent autoimmune gastric metaplasia is accompanied by expansion of epithelial populations expressing cancer progenitor cell markers. Gastric tissues from Rag°MHCII^+^, Rag°MHCII°, and Rag°MHCII° hosts co-transferred with Rag° bone marrow (BM) was enzymatically digested following adoptive transfer of CTLA4KD donor splenocytes. Non-hematopoietic cells were identified by gating on singlets, live cells, and CD45^-^ events and subjected to 40-marker spectral flow cytometry. **(A)** SPADE (Spanning-tree Progression Analysis of Density-normalized Events) trees depicting the non-hematopoietic gastric compartment from Rag°MHCII^+^ (left), Rag°MHCII° (center), and Rag°MHCII° + BM (right) hosts. Node color reflects MHCII signal intensity; node size reflects relative cell abundance. **(B)** Representative flow cytometry plots showing gating of four stromal populations within the CD45^-^compartment: fibroblastic reticular cells (FRC; PDPN^+^CD31^-^), lymphatic endothelial cells (LEC; PDPN^+^CD31^+^), blood endothelial cells (BEC; PDPN^-^CD31^+^), and double-negative stromal cells (DN; PDPN^-^CD31^-^). **(C)** Summary percentage of each stromal population across experimental groups. **(D)** Representative flow cytometry plots of MHCII expression on DN EpCAM^+^ gastric epithelial cells. **(E)** Representative flow cytometry plots showing co-expression of Sca-1 and DMBT1 within the DN EpCAM^+^ gastric epithelial population. **(F)** Summary percentage of the frequencies of Sca-1^+^DMBT1^+^, Sca-1^+^DMBT1^-^, and Sca-1^-^DMBT1^+^ within the epithelial compartment. **(G)** Summary percentage of total Sca-1^+^ and total DMBT1^+^ cell frequencies within the gastric epithelial compartment across all groups. **(H)** Representative flow cytometry plots of CD133 and PD-L1 co-expression within the DN EpCAM^+^ gastric epithelial population. **(I)** Summary quantification of CD133^+^ and PD-L1^+^ cell frequencies within the gastric epithelial compartment. Each data point represents an individual mouse; (mean ± SEM; Mann-Whitney test), \**p* < 0.05, \*\**p* < 0.01, \*\*\**p* < 0.001.

Within the CD45^-^ compartment, we identified four stromal subsets: fibroblastic reticular cells (FRC), lymphatic endothelial cells (LEC), blood endothelial cells (BEC), and double-negative cells (DN), using established markers defined in the mesenteric lymph nodes^19^ (Fig. 2B, Suppl. Fig.1). BEC frequency was significantly increased in Rag°MHCII^+^ recipients, suggesting MHCII-dependent inflammation promotes gastric angiogenesis (Fig. 2C). MHCII expression was detected across all four stromal subsets, indicating that MHCII expression is not restricted to gastric epithelial cells (Suppl. Fig. 2A, B). Gating on DN EpCAM^+^ gastric epithelial cells confirmed robust MHCII expression in Rag°MHCII^+^ hosts and its absence in Rag°MHCII° hosts (Fig. 2D).

To determine whether epithelial MHCII activation was associated with expression of cancer progenitor features, we examined expression of Sca-1 (Ly6A), a glycoprotein associated with cancer stem-like cell populations in murine malignancies and Deleted in Malignant Brain Tumor 1 (DMBT1), a putative tumor suppressor gene, within the gastric epithelial compartment. Sca-1^+^DMBT1^+^ and Sca-1^+^DMBT1^-^ cell frequencies were significantly increased in Rag°MHCII^+^ recipients relative to both MHCII KO and KO+BM (Fig. 2E, F). Total Sca-1^+^ cell frequency was significantly elevated in Rag°MHCII^+^ hosts, whereas total DMBT1^+^ frequency was unchanged, suggesting DMBT1 upregulation is restricted to the Sca-1^+^ progenitor-like subset (Fig. 2G). Although Sca-1 lacks a direct human ortholog, expansion of Sca-1^+^ cells within the epithelial compartment in MHCII-sufficient hosts suggests that epithelial MHCII activation promotes acquisition of progenitor-associated features within the gastric epithelium. Consistent with these findings, the frequency of CD133^+^PDL1^+^ epithelial cells was significantly increased in Rag°MHCII^+^ recipients compared to Rag°MHCII° and Rag°MHCII° + BM (Fig. 2H, I). Collectively, these data demonstrate that epithelial MHCII-dependent inflammation drives expansion of a gastric epithelial subpopulation co-expressing cancer progenitor and immune checkpoint markers, consistent with early premalignant transformation.

### Epithelial MHCII promotes innate and adaptive immune activation in gastric mucosa during inflammatory tumorigenesis

To characterize the hematopoietic gastric compartment, we performed SPADE clustering of CD45^+^ cells from all three experimental groups. MHCII signal intensity was enriched within the natural killer cell (NK1.1) cluster of Rag°MHCII^+^ hosts, diminished in Rag°MHCII° hosts, and restored in Rag°MHCII° + BM hosts, confirming reconstitution of MHCII-expressing innate immune populations (Fig. 3A). Notably, restoration of hematopoietic MHCII in Rag°MHCII° + BM hosts was not sufficient to rescue SPEM, further implicating epithelial rather than hematopoietic MHCII as the driver of premalignant transformation.

**Figure 3.**
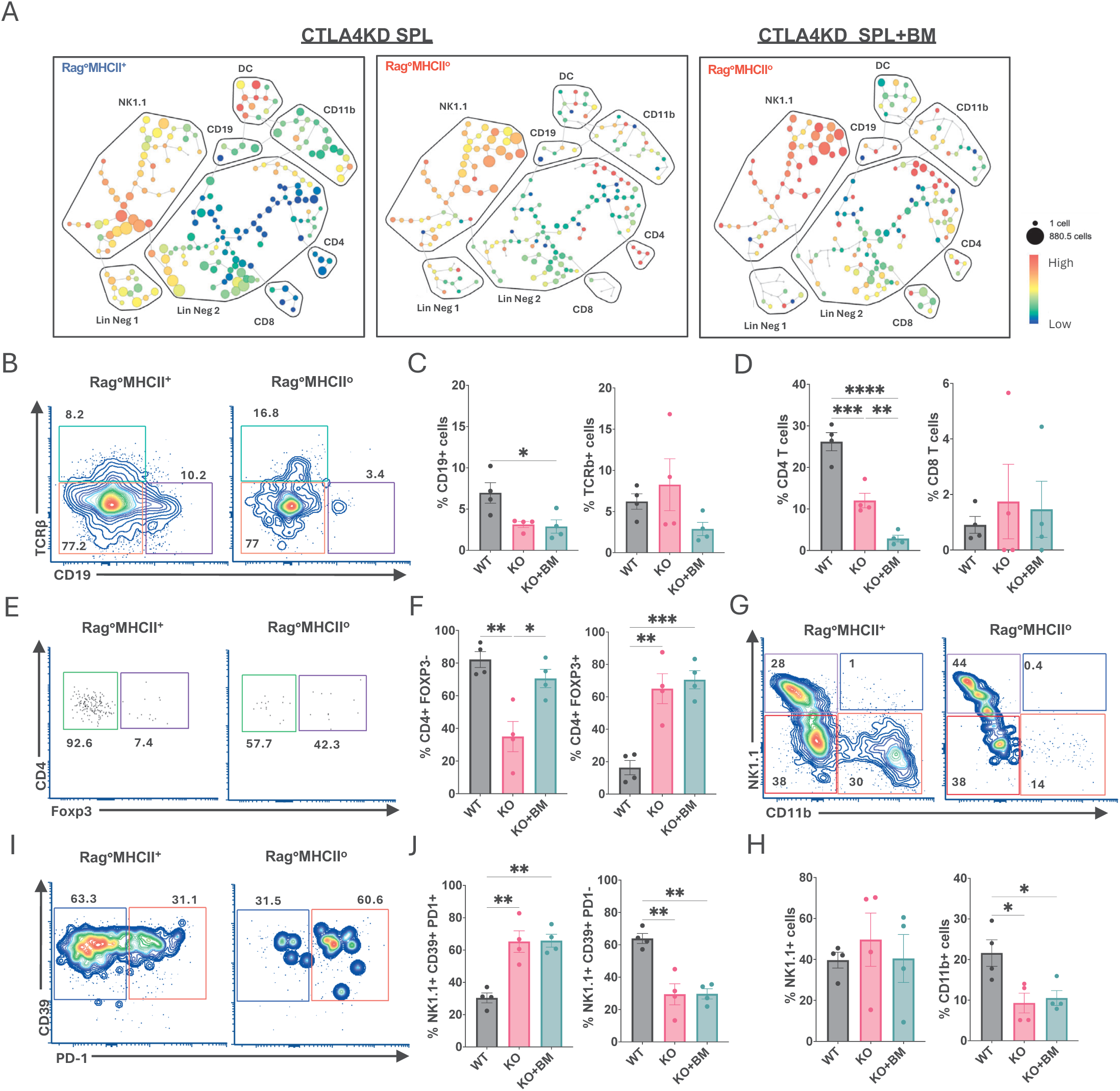
Epithelial MHCII expression promotes expansion of adaptive and innate immune cells into the gastric mucosa during inflammatory tumorigenesis. Gastric tissues from Rag°MHCII^+^, Rag°MHCII°, and Rag°MHCII° hosts co-transferred with Rag° bone marrow (BM) was enzymatically digested following adoptive transfer of CTLA4KD donor splenocytes. Hematopoietic cells were identified by gating on singlets, live cells, and CD45^+^ and subjected to 40-marker spectral flow cytometry. **(A)** SPADE (Spanning-tree Progression Analysis of Density-normalized Events) trees depicting the hematopoietic compartment in gastric tissue from Rag°MHCII^+^ (left), Rag°MHCII° (center), and Rag°MHCII° + BM (right) hosts. Node color reflects MHCII signal intensity; node size reflects relative cell abundance. **(B)** Representative flow cytometry plots showing CD19 (B cells) and TCRβ (T cells) expression in the gastric CD45^+^ compartment. **(C)** Summary percentage of CD19^+^ and TCRβ^+^ cell frequencies. **(D)** Summary percentage of CD4^+^ and CD8^+^ T cell frequencies, gated on TCRβ^+^. **(E)** Representative flow cytometry plots of CD4 and Foxp3 expression within the gastric CD45^+^ compartment. **(F)** Summary percentage of CD4^+^Foxp3^+^ regulatory T cells (T_regs_) and CD4^+^Foxp3^-^ conventional T cells (T_conv_) frequencies. **(G)** Representative flow cytometry plots of NK1.1 and CD11b expression, gated on TCRβ^-^CD19^-^. **(H)** Summary percentage of NK1.1^+^ and CD11b^+^ cell frequencies across groups. **(I)** Representative flow cytometry plots of PD-1 and CD39 expression within the NK1.1^+^ cell gate. **(J)** Summary percentage of PD-1^+^ and CD39^+^ frequencies within the NK1.1^+^ population across groups. Each data point represents an individual mouse (mean ± SEM; Mann-Whitney test), \**p* < 0.05, \*\**p* < 0.01, \*\*\**p* < 0.001.

Further analysis revealed an increased frequency of CD19^+^ B cells in Rag°MHCII°, with no significant difference in total TCRβ^+^ T cell frequency (Fig. 3B, C). CD8^+^ T cell frequency did not differ between groups, whereas CD4^+^ T cell frequency was significantly elevated in Rag°MHCII^+^ (Fig. 3D). Effector CD4^+^ T cells (CD4^+^Foxp3^-^) were significantly expanded in Rag°MHCII^+^, reduced in Rag°MHCII°, and recovered in Rag°MHCII° + BM recipients despite absent SPEM pathology, indicating the immune cell MHCII supports CD4 expansion but cannot substitute epithelial MHCII in driving SPEM. Conversely, CD4^+^Foxp3^+^ regulatory T cell (T_reg_) frequency was significantly reduced in Rag°MHCII°, suggesting this inflammatory environment favors effector over T_reg_ CD4 differentiation (Fig. 3E, F).

Analysis of innate immune cells revealed a decreased frequency of myeloid CD11b^+^ cells in Rag°MHCII°, indicating that in this context, inflammation promotes myeloid cell accumulation in the gastric mucosa, and that recruitment is decreased in the absence of epithelial MHCII (Fig. 3G, H). Natural killer cell (NK) frequency did not differ significantly between groups, however NK cells in Rag°MHCII^+^ mice were enriched for a CD39^+^PD1^-^ activated phenotype, whereas both KO and KO+BM groups expressed an expanded CD39^+^PD1^+^ exhaustion-associated phenotype (Fig. 3I, J). Taken together, these results demonstrate that epithelial MHCII orchestrates a chronic inflammatory immune environment in the gastric mucosa, characterized by effector CD4 expansion, T_reg_ suppression, myeloid accumulation, and NK cell activation, establishing the immune conditions required for epithelial reprogramming and premalignant transformation.

### Systemic immune effects during autoimmune-driven gastric tumorigenesis

To determine whether immune modulation driven by gastric inflammation extends beyond the gastric mucosa, we examined the splenic immune compartment of Rag°MHCII^+^, Rag°MHCII°, and Rag°MHCII° + BM mice. CD19^+^ B cell frequency was significantly elevated in Rag°MHCII^+^, while total TCRβ^+^ T cell frequency did not differ between groups (Fig. 4A, B). Similar to the gastric tissue, CD4^+^ T cell frequency was significantly increased in Rag°MHCII^+^ with no changes observed across CD8^+^ T populations (Fig. 4C, D). To evaluate the CD4 T cell subsets, we examined Gata3 and Foxp3, the master transcription factors for Th2 and Treg differentiation, respectively. Strikingly, Gata3^+^Foxp3^-^ Th2-like effector CD4 cells were expanded in Rag°MHCII^+^ in comparison to both Rag°MHCII° and Rag°MHCII° + BM, but no differences were seen in the Gata3^+^Foxp3^+^ population across cohorts (Fig. 4E, F). Further analysis revealed an elevated frequency of Ki67^+^PD1^-^ CD4 cells in Rag°MHCII^+^ mice, while Ki67^-^PD1^+^ cells did not differ between groups, suggesting increased proliferation of CD4 cells in the Rag°MHCII^+^ cohort (Fig. 4G, H). Together, these findings demonstrate that epithelial MHCII is required for systemic CD4 T cell expansion, Th2 polarization, and effective proliferation, and that its absence results in a systemic quiescent peripheral CD4 compartment despite the presence of MHCII^+^ APCs in the Rag°MHCII° + BM mice.

**Figure 4.**
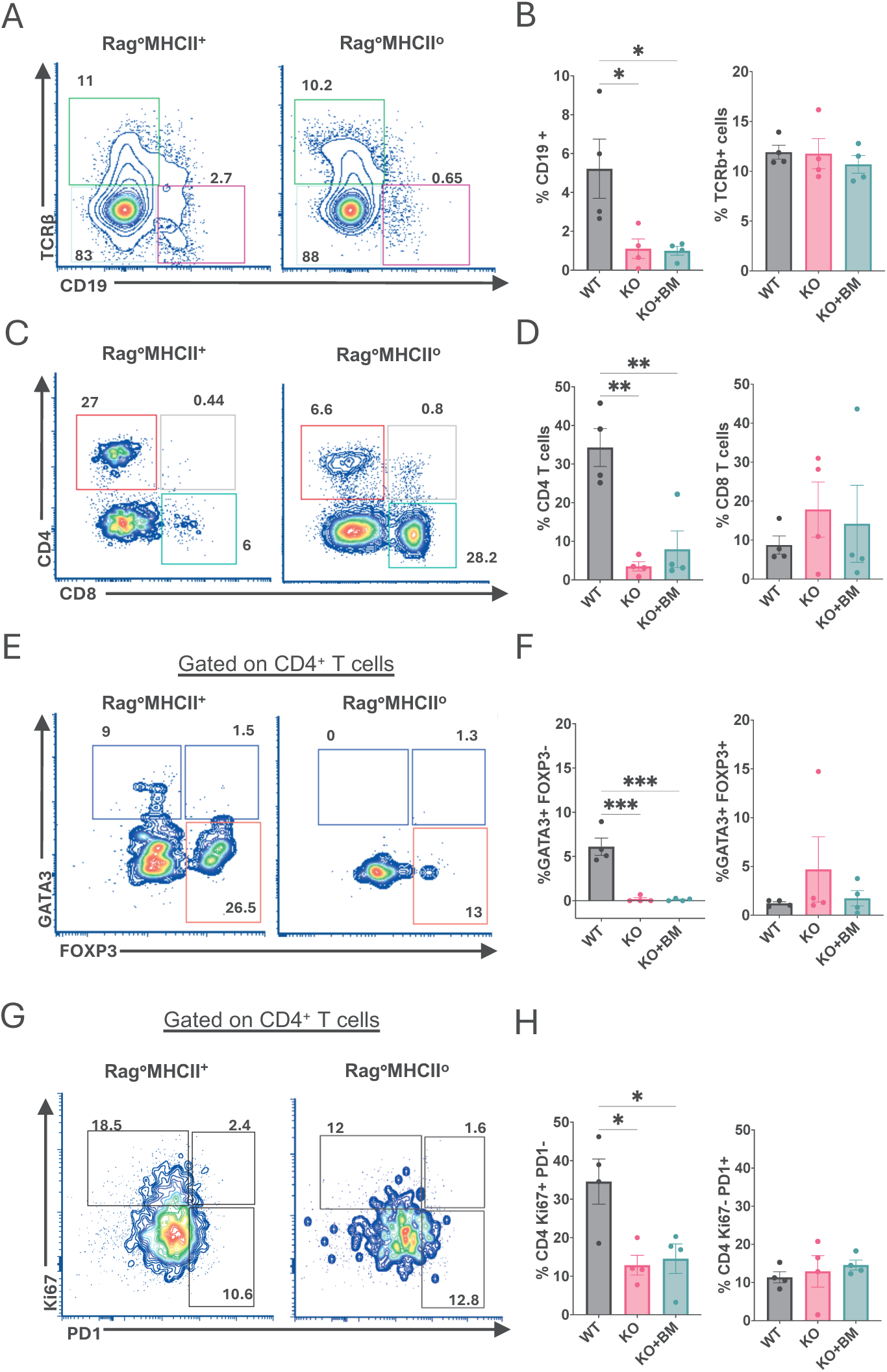
Systemic immune effects during autoimmunity-driven gastric tumorigenesis. Spleens Rag°MHCII^+^, Rag°MHCII°, and Rag°MHCII° hosts co-transferred with Rag° bone marrow (BM) was harvested and analyzed by flow cytometry following CTLA4KD adoptive transfer. **(A)** Representative flow cytometry plots showing CD19 (B cells) and TCRβ (T cells) expression. **(B)** Summary percentage of CD19^+^ and TCRβ^+^ cell frequencies. **(C)** Representative flow cytometry plots showing CD4, and CD8 T cells expression gated on TCRβ^+^ splenic T cells. **(D)** Summary percentage of CD4^+^ and CD8^+^ T cell frequencies. **(E)** Representative flow cytometry plots of Foxp3 and Gata3 expression, gated on splenic CD4^+^ T cells. **(F)** Summary percentage of Gata3^+^Foxp3^-^ and Gata3^+^Foxp3^+^ CD4 T subset frequencies. **(G)** Representative flow cytometry plots of Ki67 and PD1 expression on splenic CD4^+^ T cells. **(H)** Summary percentage of Ki67^+^PD1^-^and Ki67^-^PD1^+^ cell frequencies across groups. Each data point represents an individual mouse (mean ± SEM; Mann-Whitney test), \**p* < 0.05, \*\**p* < 0.01, \*\*\**p* < 0.001.

### MHCII colocalizes with EpCAM in gastric tissue during SPEM

To directly visualize epithelial MHCII expression and CD4^+^ T cell distribution within the gastric mucosal architecture, we performed immunofluorescence on gastric tissue sections (Fig. 5). EpCAM and MHCII signals showed significant colocalization in the stomach of Rag°MHCII^+^ but not in Rag°MHCII° (Fig. 5A, B). We applied automated cell classification using QuPath (Suppl. Fig. 3A, B) and identified CD4^+^ T cells in proximity to EpCAM^+^MHCII^+^ glands in Rag°MHCII^+^, with a mean nearest neighbor distance of 29 to 115 µm across animals, and a subset observed in direct contact with glands expressing MHCII (Fig. 5C, D). QuPath analysis of Rag°MHCII° tissue confirmed the absence of EpCAM^+^MHCII^+^ cells (Suppl. Fig. 3B). Notably, CD4^+^ T cell density was significantly higher in Rag°MHCII^+^ compared to Rag°MHCII° hosts, consistent with the flow cytometric data (Fig. 5D).

**Figure 5.**
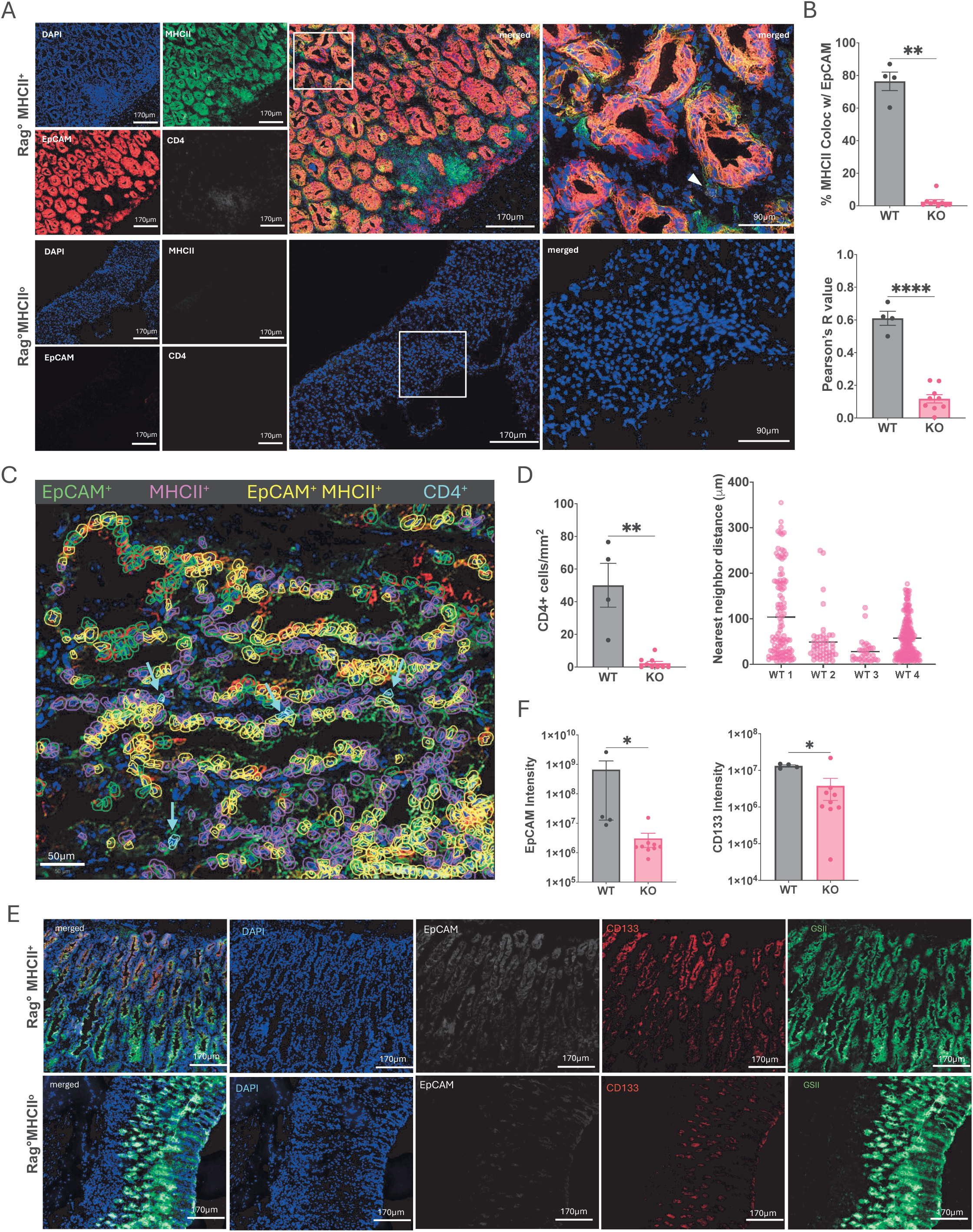
Colocalization of MHCII and EpCAM epithelial cells during gastric metaplasia. **(A)** Representative immunofluorescence images of stomach tissue from Rag°MHCII^+^ and Rag°MHCII° post CTLA4KD adoptive transfer, stained for EpCAM (red), MHCII (green), CD4 (white), and DAPI (blue) **(B)** Colocalization analysis and Pearson correlation coefficient R values of EpCAM and MHCII signals using the Coloc2 plugin in Fiji/ImageJ. **(C)** Representative immunofluorescence image of gastric tissue from a Rag°MHCII^+^ host with automated QuPath cell classification overlay, excluding unclassified cells. EpCAM^+^MHCII^+^ epithelial cells (yellow), CD4^+^ T cells (cyan), EpCAM^+^ epithelial cells (green), and MHCII^+^ cells (magenta) are shown. Image represents a magnified region of interest (ROI). Scale bar, 50µm. **(D)** Average CD4^+^ cells quantified as cells per mm2 of annotated gastric tissue ROI in Rag°MHCII^+^ and Rag°MHCII° hosts (left). Each point represents one mouse, mean ± SEM; Mann-Whitney test. **p < 0.01. Nearest neighbor distance from each CD4^+^ cell to the closest EpCAM^+^MHCII^+^ epithelial cell in Rag°MHCII^+^hosts in µm (right). Each point represents an individual CD4^+^. **(E)** Representative immunofluorescence images of gastric tissue sections from Rag°MHCII^+^ and Rag°MHCII° stained for EpCAM (white), CD133 (red), GSII (green), and DAPI (blue). **(F)** Fluorescence intensity quantification of indicated markers within defined tissue ROIs. Corrected total cell fluorescence (CTCF) was calculated as: integrated density - (ROI area x mean background fluorescence), measured from two to five fields per tissue and averaged per mouse. Each data point represents one mouse; mean ± SEM; Mann-Whitney test. *p < 0.05, **p < 0.01, ***p < 0.001.

To detect metaplastic lesions in the gastric mucosa, we stained for established markers of SPEM, and markers identified through flow cytometry analysis. EpCAM and CD133 signal intensities were significantly elevated in Rag°MHCII^+^ relative to Rag°MHCII° mice, consistent with the expansion of EpCAM^+^ epithelial cells and CD133^+^ progenitor populations identified by flow cytometry (Fig. 5E, F). GSII lectin staining, which marks mucous neck cells and metaplastic epithelium, revealed a striking spatial redistribution in Rag°MHCII^+^ mice, shifting from expression restricted to mucous neck cells at the luminal surface in Rag°MHCII° tissue to a diffuse glandular distribution throughout the gastric mucosa (Fig. 5E). Total GSII signal intensity did not differ significantly between groups (Suppl. Fig. 3C), indicating that SPEM-associated changes in GSII localization reflect epithelial reprogramming and glandular redistribution.

### Early-stage gastric premalignancy can be reversed by removal of epithelial MHCII

We next asked whether ongoing epithelial MHCII expression is required to sustain premalignant transformation once it has already been initiated. To address this while controlling for MHCII contributions from transferred immune cells, CTLA4KD splenocytes were depleted of CD19^+^, CD11c^+^, CD11b^+^, NK1.1^+^, Gr1^+^ and F4/80^+^ cells prior to adoptive transfer, achieving greater than 99% depletion efficiency (Suppl. Fig. 4A). Depleted splenocytes were transferred into Rag°H2Ab^fl/fl^Rosa26-CreERT2 neonatal mice, where tamoxifen treatment induced Cre-mediated deletion of H2Ab with 99.6-99.8% efficiency (Suppl. Fig. 4B). At 3 months post-transfer, coinciding with early SPEM development, recipients received tamoxifen or vehicle injections. Tissue was harvested for analysis 2 weeks post-tamoxifen or post-vehicle treatment (Fig. 6A). Strikingly, tamoxifen-treated mice showed no histological evidence of SPEM, whereas vehicle-treated hosts developed robust premalignant pathology comparable to Rag°MHCII^+^ mice (Fig. 6B). Histopathological scoring confirmed significantly elevated gastric inflammation and SPEM severity in vehicle-treated mice relative to tamoxifen-treated mice (Fig. 6C).

**Figure 6.**
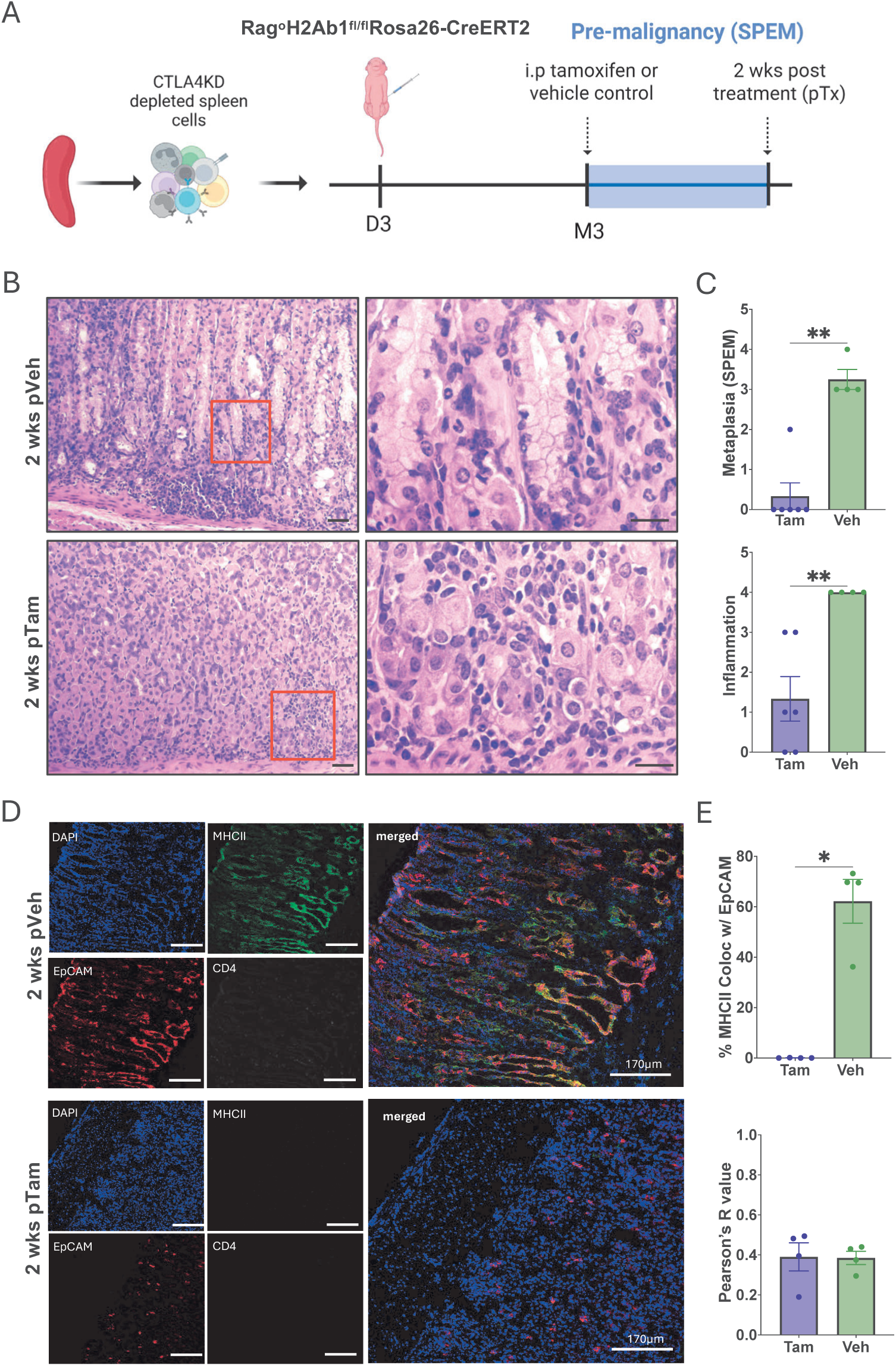
Early-stage gastric premalignancy can be reversed by removing epithelial MHCII. **(A)** Experimental schematic. Spleen cells isolated from CLA4KD donor mice were depleted of NK1.1, CD11b, CD11c, F4/80, Gr1 and CD19. Depleted cells adoptively transferred into Rag°H2Ab^fL/fL^Rosa26-CreERT2 neonates (3-5 days old). Intraperitoneal (i.p) tamoxifen or vehicle control was given during SPEM development at 3 months (M3) post adoptive transfer. Animals were analyzed between 2-3 weeks post treatment (pTX) with tamoxifen or vehicle control. **(B)** Representative H&E-stained gastric sections from tamoxifen or vehicle treated hosts at 4X (left), and 40X (right). **(C)** Histopathology scores for gastric inflammation and SPEM from cohorts described in (B). **(D)** Immunofluorescence images of gastric tissue sections from tamoxifen and vehicle treated controls, stained for EpCAM (red), MHCII (green), CD4 (white), and DAPI (blue) **(E)** Colocalization analysis, and Pearson correlation coefficient R values of EpCAM and MHCII signals using the Coloc2 plugin in Fiji/ImageJ. H&E Scale bar, 4X:100 µm, and 40X: 50 µm. Each point represents one mouse; mean ± SEM; Mann-Whitney test. *p < 0.05, **p < 0.01, ***p < 0.001.

Immunofluorescence analysis confirmed that vehicle-treated hosts exhibited significant EpCAM-MHCII colocalization within the gastric epithelial compartment, consistent with intact epithelial MHCII expression, whereas tamoxifen-treated hosts showed markedly reduced colocalization. Pearson R values did not differ significantly between groups, despite a significant decrease in colocalization, indicating that the loss of MHCII-EpCAM overlap is spatially restricted and does not alter overall signal relationship (Fig. 6D, E). Collectively, these data demonstrate that epithelial MHCII is required not only to initiate but to sustain premalignant gastric transformation. Removal of MHCII after CD4 T cell engraftment and early disease onset is sufficient to abolish SPEM development, identifying ongoing epithelial antigen presentation as a targetable checkpoint in autoimmunity-driven gastric premalignancy.

### Single-cell transcriptomic analysis identifies MHCII-mediated signaling from SPEM to T cells

To identify molecular interactions underlying epithelial MHCII and immune activation at the transcriptomic level, we analyzed scRNAseq data from CTLA4KD gastric tissue at premalignant and adenocarcinoma stages, and age-matched controls. Unsupervised clustering identified a metaplastic/tumorigenic epithelial cluster defined by expression of SPEM-associated genes, Gkn3, Muc6, and Aqp5. A T/Mast cell cluster was also defined by expression of Ptprc, CD52, Rgs1, and Coro1a. CellChat ligand-receptor interaction analysis predicted significant MHCII pathway signaling from the Metaplastic/Tumorigenic cluster to the T cell cluster at the premalignant stage (Fig. 7A). Consistent with this predicted interaction, the MHCII ligand H2-Ab1 was enriched in the Metaplastic/Tumorigenic cluster while the CD4 receptor was restricted to the T cell cluster across both disease stages (Fig. 7B). These data provide independent transcriptomic corroboration of the genetic requirement for epithelial MHCII in CD4 T cell-driven premalignant gastric transformation.

**Figure 7.**
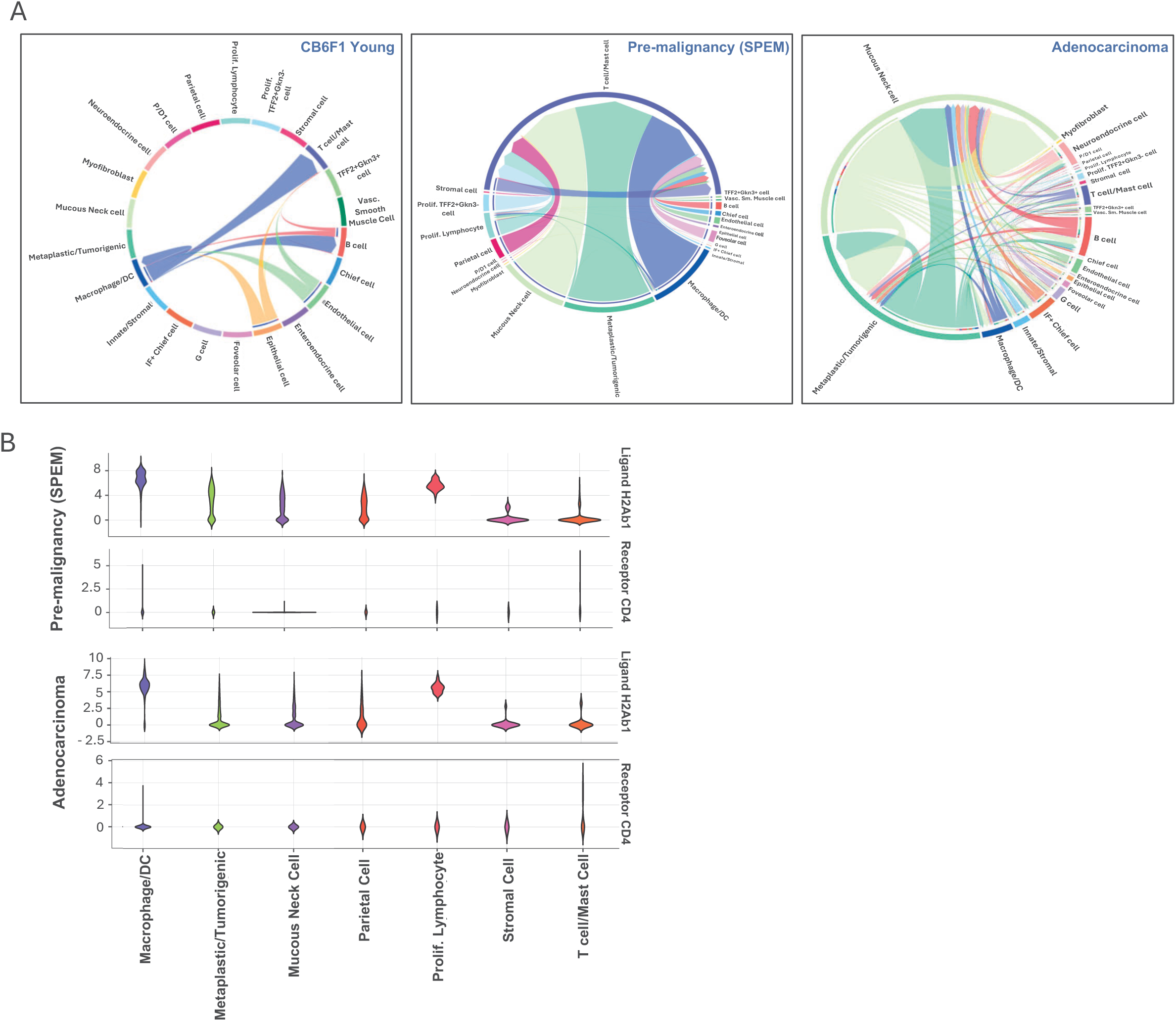
Single-cell transcriptomic analysis identifies MHCII-mediated signaling from metaplastic cells to T cells in gastric tumorigenesis. Gastric mucosa of age-matched CB6F1 and CTLA4KD mice were enzymatically digested and analyzed by single cell RNA sequencing (scRNAseq) analysis. Clustering of cell populations was conducted using the Serat package on R. **(A)** Cell-cell communication analysis using CellChat identifying MHCII signaling pathways in CB6F1 young (left), CTLA4KD young (center) and CTLA4KD old (right). Chord diagrams depict predicted ligand-receptor interactions; arrow directionality indicates sending and receiving populations. **(B)** Violin plots showing expression of H2-Ab1 and CD4 across selected cell clusters in CTLA4KD young (top) and CTLA4KD old (bottom).

### MHCII expression in human gastric adenocarcinoma

To evaluate the translational relevance of our findings in human disease, we obtained human gastric adenocarcinoma tissue and matched healthy gastric tissue. This dataset consists of specimens from 11 patients, both females and males, ranging in age 22 to 73. We performed immunofluorescence staining and applied automated cell classification using QuPath to quantify population densities across the cohort (Fig. 8A, Suppl. Fig. 5A). EpCAM^+^MHCII^+^ epithelial cell density was significantly higher in gastric adenocarcinoma tissue compared to matched healthy gastric mucosa from the same patient (Fig. 8B), directly recapitulating epithelial MHCII upregulation identified in the CTLA4KD mouse model. Total EpCAM^+^ epithelial cell density was also significantly elevated in adenocarcinoma relative to matched healthy tissue (Fig. 8B), consistent with the epithelial expansion characteristic of gastric adenocarcinoma and confirming that the QuPath classification pipeline detected biologically relevant differences between tissue types. MHCII^+^ and GSII^+^ metaplastic cell density did not differ significantly between adenocarcinoma and matched healthy tissue (Fig. S5, B). Collectively, these findings demonstrate that epithelial MHCII upregulation is a feature of human gastric adenocarcinoma.

**Figure 8.**
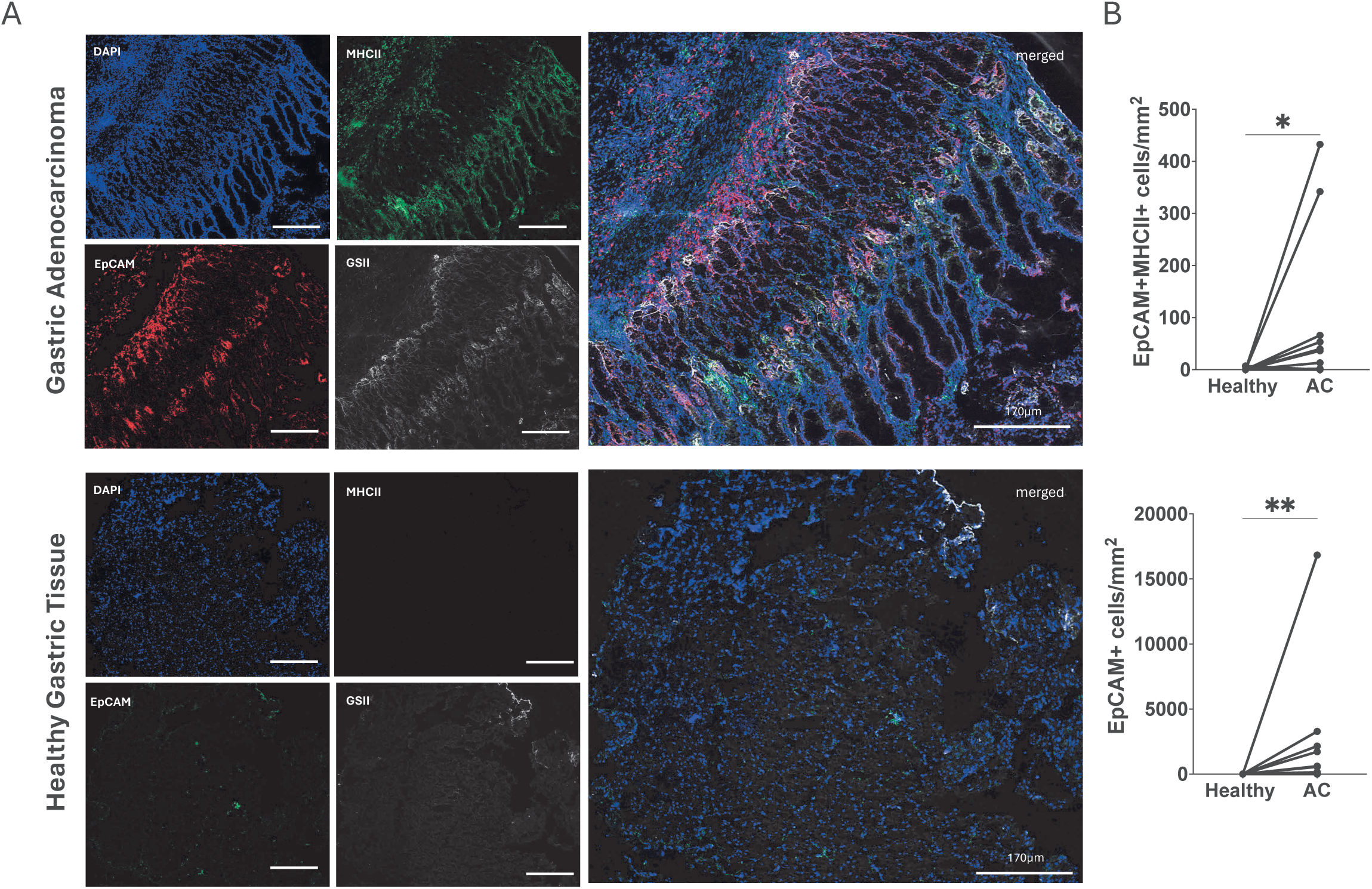
Detection of MHCII expression on epithelial cells in human gastric adenocarcinoma. **(A)** Representative immunofluorescence images of human matched gastric adenocarcinoma (top) and healthy tissue (bottom), stained for EpCAM (red), MHCII (green), GSII (white), and DAPI (blue). **(B)** EpCAM^+^MHCII^+^ (top) and EpCAM^+^ (bottom) cells quantified as cells per mm2 of annotated gastric tissue ROI in human adenocarcinoma and matched healthy controls. Each point represents one patient, n=11; Wilcoxon matched pairs signed rank tests; *p < 0.05, **p < 0.01.

## Discussion

Chronic autoimmune inflammation is an established driver of epithelial cancer risk across multiple organ systems, from autoimmune gastritis to inflammatory bowel disease and primary biliary cholangitis^20–23^. Yet much remains to be learned about the molecular mechanism that connects autoreactive immune activation to premalignant epithelial transformation. Here, we demonstrate that gastric epithelial cells are active participants in autoimmunity-driven premalignancy, expressing functional MHC class II (MHCII) that is required for CD4 T cell-mediated premalignant transformation in a manner that professional antigen-presenting cells cannot replicate.

Expression of MHCII has long been considered a defining feature of professional antigen-presenting cells (APCs), yet non-hematopoietic cells, including intestinal and airway epithelial cells, have been shown to express MHCII under inflammatory conditions^24,25^. In our model, we identified that non-hematopoietic MHCII drives the initiation of gastric tumorigenesis. The failure of BM reconstitution in reestablishing SPEM pathology, despite restoring MHCII expression in BM-derived immune cells, demonstrates that MHCII on non-hematopoietic cells plays a crucial role in driving premalignancy. It is worth noting that initiation of gastric tumorigenesis was dependent on specific antigen presentation. Disruption of antigenic presentation by knocking out H2-DM resulted in a complete loss of pathology.

In addition to driving SPEM pathology, we uncovered that autoimmunity-driven inflammation was associated with the expansion of cancer progenitor and immune-evasive epithelial populations. We observed upregulation of Sca-1 (Ly6A) and CD133+PDL1+ in gastric epithelial cells of MHCII-sufficient hosts. Sca-1 and CD133 (prominin-1) are cancer stem cell markers associated with enhanced tumor initiation, migration, and invasion^26–30^, while PD-L1 has been extensively characterized as a mediator of tumor immune suppression through engagement of PD-1 on T cells ^12,13^. The expansion of these populations, specifically in the context of functional epithelial MHCII expression, suggests that the MHCII-dependent inflammatory environment does not simply cause generalized epithelial stress but actively shapes the premalignant epithelial phenotype toward a progenitor-like and immune-evasive state. Most striking is the co-expression of MHCII and PD-L1 on the same epithelial cells, which can activate CD4 T cells through antigen presentation and suppress T cell effector function through PD-1 engagement, suggesting that the premalignant epithelium acquires immune-evasive features early in the transformation process.

Previous work from our laboratory established that CD4^+^CD25^-^ T cells and IL-4/IL-13 are required for SPEM development in the CTLA4KD model, with CD4 Th2 effector cells identified as the likely pathogenic driver^9^. In the gastric mucosa, we observed a complex chronic inflammatory process, characterized by increased frequency of infiltrating immune cells from both the innate and adaptive branches. Notably, frequencies of CD19^+^ and CD4^+^ cells increased in MHCII-sufficient hosts. Further characterization of CD4 T cells revealed that BM-reconstitution was able to restore the CD4^+^Foxp3^-^ T conventional cells to frequencies seen on MHCII-sufficient host, while also maintaining a high CD4^+^Foxp3^+^ T_regs_ observed on the KO and KO+BM mice. This could indicate that while BM-derived MHCII can restore CD4 T_conv_ to wildtype levels and maintain a local immunosuppressive environment through increased frequencies of Tregs, it is not sufficient to drive autoimmune tumorigenesis, likely requiring specific antigen presentation by epithelial cells. Within the innate immune compartment, autoimmunity-driven inflammation was associated with enrichment of CD11b^+^ myeloid cells in the gastric mucosa, consistent with the known capacity of Th2 cytokines to promote myeloid recruitment at sites of chronic mucosal inflammation^31,32^, and with a shift in NK cell activation states^33,34^. Whether these innate immune changes contribute mechanistically to premalignant transformation or reflect the broader inflammatory state downstream of the CD4 Th2 response remains to be determined.

Beyond the gastric mucosa, autoimmunity-driven tumorigenesis presents a striking systemic effect on modulating adaptive immunity. Notably, we observed a robust expansion of Gata3^+^Foxp3^-^ Th2 CD4 cells in MHCII WT mice, directly linking the Th2 polarization required for SPEM development as previously described^9^. Additionally, Ki67^+^PD1^-^ CD4 T cells were significantly enriched in MHCII WT mice during premalignancy, suggesting that the autoimmune inflammatory environment supports productive CD4 proliferation systemically. Together, these systemic findings suggest that the autoimmunity-driven inflammation extends beyond the gastric mucosa to systemically shape the state and polarization of the peripheral CD4 response.

It is worth noting that premalignancy represents a critical window of opportunity for cancer prevention^35,36^. Understanding the signals that initiate and maintain epithelial tissue remodeling during tumorigenesis is essential for the implementation of therapeutic strategies. Single-cell RNA sequencing predicted MHCII pathway signaling between the metaplastic/tumorigenic cluster to the T cell clusters. Communication probability was most prominent in the premalignant stage, highlighting premalignancy as a key stage of epithelial-immune cross-talk during cancer development. Importantly, we identified epithelial MHCII, not only as a player in initiation of autoimmunity-driven tumorigenesis, but also in maintaining cancer progression. Indeed, removing epithelial MHCII after the establishment of premalignancy in the gastric tissue, reversed SPEM development in mice.

Emerging evidence suggests cancer cells utilize MHCII to evade immune responses, but the mechanism remains unclear. A recent study showed that a subset of MHCII-expressing breast cancer cells lacking co-stimulatory molecules promoted tumor metastasis and immune evasion^37^. Additionally, single-cell analysis in pancreatic cancer identified cancer-associated fibroblasts with high levels of MHCII but also lacking costimulatory molecules^38^. Similar to these observations, we identified increased EpCAM^+^MHCII^+^ cells in human adenocarcinoma tissue samples, relative to matched healthy controls. While host factors such as *H. pylori* status and patients’ genetic risk factors can influence tumor heterogeneity, the functional significance of MHCII on cancer cells is context-dependent. For example, tumor specific MHCII was associated with improved survival in melanoma and classic Hodgkin lymphoma patients treated with anti-PD-1/anti-PD-L1 but was associated with resistance to anti-CTLA4 immunotherapies^39–41^. However, epithelial MHCII in human gastric cancer may stratify patients by immune microenvironment, with patients with high epithelial MHCII having distinct CD4 T cell and immune checkpoint therapy responses.

Overall, our findings provide insight into how epithelial MHCII plays a critical role in initiating autoimmunity-driven tumorigenesis and sustaining premalignancy growth in the stomach. Collectively, our data shows MHCII expression on epithelial cells, serves as a main driver of epithelial differentiation as well as systemic and local immune modulation. These findings raise the question of whether epithelial MHCII plays a similar role in other autoimmunity-driven cancers, such as Barrett’s esophagus, colitis-associated colorectal cancer, and cholangiocarcinoma. While CTLA4 haploinsufficiency is rare, the CTLA4KD mice capture a unifying point in which chronic inflammation, whether from *H. Pylori* or immune dysregulation, leads to progression from premalignancy (SPEM) to cancer (GA). The premalignant window is one of the most clinically actionable points for intervention. If epithelial MHCII is required to sustain premalignancy, targeting the epithelial MHCII-CD4 interface may represent a therapeutic strategy for high-risk patients. Future work should be aimed at identifying the specific epithelial peptides presented by gastric epithelial MHCII, as well as the downstream transcriptional response in the gastric epithelium.

## Methods

### Mice

The following transgenic and knockout mice were used in this study: RAG1 knockout (Rag°), Rag-deficient H2DM-knockout (Rag°H2DM°), Rag-deficient MHCII-knockout (Rag°MHCII°), Rag-deficient MHCII-wildtype (Rag°MHCII^+^), Rag°H2Ab1^fl/fl^Rosa26-CreERT2 (triple knockout mice, conditional MHCII deletion). Mice were on the C57BL/6 (B6) genetic background. These mutant strains were obtained from The Jackson Laboratory (Bar Harbor, ME). CTLA4KD were on the Balb/c x C57BL/6 F1 background and previously described^9^. Animals were maintained in a specific pathogen-free facility. In our experimental system, we did not observe a difference between male and female animals. Therefore, both sexes with littermate controls were used in this study. The studies were approved by the Institutional Animal Care and Use Committee at the University of Miami.

### Mucosal cell isolation

Mouse gastric mucosa was excised and cut into pieces <5mm in size. Tissue was incubated in RPMI media (#22400-089, Gibco) supplemented with FBS (#A5256701, Gibco) and pen/step (#15140-122, Gibco), containing 1mg/ml Collagenase IV (#C5138, Sigma-Aldrich) and 10 U/ml DNase I (#LS002007, Worthington) for 20 minutes at 37°C on a shaker. Following incubation, supernatant was collected into cell collection media (PBS+2% FBS, + 2mM Ethylenediaminetetraacetic acid) and additional digestion media was added to the stomach tissues. This was repeated twice, for a total of three 20-minute incubations. Digested tissue was mechanically dissociated using 18G needle, filtered using 70µm filter and washed with PBS (#14190-144, Gibco).

### Isolation and adoptive transfer of immune cells

Sterile single cell suspensions were made with pooled spleens from CTLA4KD donors. Red blood cells were removed by ACK lysis buffer. Each recipient was reconstituted with approximately 10 x 10^6^ cells at 2-5 days of age. Bone marrow (BM) was flushed from the femur, tibia, humerus, and ulna of Rag° mice. Recipients reconstituted with the mixture of splenocytes, and BM received a total of 13 x 10^6^ cells. In some experiments splenocytes were depleted of CD11b, CD11c, CD19, F4/80, NK1.1, and GR1 cells. Each recipient was reconstituted with approximately 3 x10^6^ cells at 2-5 days of age. Single cell suspensions were injected intraperitoneally (i.p).

### Magnetic bead-based cell isolation

A single cell suspension was prepared and labeled with biotinylated-conjugated antibodies according to standard procedure. Magnetic depletion of specific immune populations with streptavidin-conjugated magnetic beads from New England Biosciences. A mixture of biotinylated antibodies including anti-CD11b, anti-CD11c, anti-CD19, anti-F4/80, anti-NK1.1 and anti-GR1 was used in preparation for adoptive transfer experiment of depleted spleen cells. Purity was assessed via flow cytometry analysis.

### Flow cytometry analysis

Single-cell preparations from the gastric mucosa were analyzed using a 40-color spectral flow cytometry panel previously described ^42^. Flow cytometry analyses were conducted using a standard procedure. Cells were blocked against nonspecific antibody binding using normal mouse serum (#SP-0002-VX5, ImmunoReagents), anti-CD16/32 antibody (Clone 2.4G2), BD Horizon Brilliant Stain Buffer Plus (3566385, BD) and CellBlox blocking buffer (#C001T02F01). After blocking, cells were stained with surface antibodies, and the fixed and permeabilized for intracellular staining using the Transcription Factor Buffer Set (#562574, BD). Dead cells were excluded using Fixable Viability Dye eFluor 780 (Thermo Fisher). Doublets were gated out from analysis. Manual gating was conducted using FCS express software. Gastric samples were analyzed using the Cytek Aurora from Cytek Biosciences; immune cells were analyzed using CytoFLEX LX Platform from Beckman Coulter.

### SPADE tree analysis

Spectral flow cytometry underwent viSNE dimensionality reducing analysis on the Cytobank platform using FCS files were pre-gated on live cells (Zombie NIR^-^) and hematopoietic (CD45^+^) cells or non-hematopoietic (CD45^-^) lineage. viSNE analysis transformed FCS data into 2 dimensions using the Barnes-Hut implementation of the t-distributed stochastic neighbor embedding (tSNE) algorithm. Analysis was performed on the samples using proportional sampling, with 7,500 iterations, a perplexity of 30, and a 8 of 0.5, following Cytobank user guides. Following viSNE analysis, unsupervised clustering was performed using the Spanning-tree Progression Analysis of Density-normalized Events (SPADE) algorithm implemented in Cytobank (Cytobank Inc.). Cells were clustered on the dimensionality reduction channels tSNE1 and tSNE2. Node size reflects relative cell abundance and node color reflects mean signal intensity for the indicated marker. Representative SPADE trees from each experimental group are shown.

### Histopathology analysis

Stomach samples were collected from euthanized animals and fixed in 10% formalin solution. Paraffin-embedded sections and H&E staining were done by the Cancer Modeling Shared Resource (CMSR) at the University of Miami Miller School of Medicine. Histopathology was documented by microscopy. Pathology scoring was performed in a blinded fashion to assess the extent of inflammatory damage and SPEM development according to criteria previously described^9^.

### Immunofluorescence microscopy

Stomachs were collected and fresh frozen in embedding media. The cryopreserved tissue blocks were sectioned with a cryostat at 6-µm thickness, fixed with acetone, and then stained with fluorescently labeled antibodies. Immunofluorescence microscopy was conducted using a standard protocol using the ECHO Revolve fluorescence microscope. The following mouse antibody conjugates were used: APC-conjugated anti-CD4 (GK1.5, eBioscience), AL488-conjugated anti-I-Ab (AF6-120.1, Biolegend), PEDazzle594-conjugated anti-EpCAM (G8.8, Biolegend), FITC-conjugated anti-EpCAM (G8.8), PEDazzle594-conjugated anti-CD133 (315-2C11, Biolegend), AF647-conjugated GSII. The following human antibody conjugates were used: PEDazzle594-conjugated anti-EpCAM (9C4, Biolegend), FITC-conjugated anti-HLA-DR-DP-DQ (TU39, Biolegend), AF647-conjugated GSII.

### Fluorescence intensity quantification and colocalization analysis

Fluorescence signal intensity and colocalization analyses were performed using Fiji/ImageJ. For corrected total cell fluorescence (CTCF) quantification, regions of interest (ROIs) were manually defined within the gastric mucosal compartment of each image. CTCF was calculated for each ROI as: CTCF = Raw Integrated Density - (ROI Area x Mean Background Fluorescence), where background fluorescence was determined as the mean value of cell-free regions. Colocalization of EpCAM and MHCII signals was quantified using the Coloc2 plugin. Manders’ Overlap Coefficients corrected for autothreshold (tM1/tM2), and Pearson correlation coefficient R, were calculated for each ROI as a measure of signal colocalization. Two to five non-overlapping fields per tissue section were quantified and averaged to yield one value per mouse.

### QuPath image analysis and spatial quantification

Immunofluorescence images were analyzed using QuPath v0.7. Individual channel images acquired at 10X magnification were merged into multi-channel composite TIFF files using Fiji/ImageJ prior to import. Pixel size was set to 0.3551 µm/pixel based on instrument specifications. Tissue regions of interest (ROIs) were manually annotated to delineate the gastric mucosa and submucosa, excluding the muscularis propria and tissue edge. Cell detection was performed on the DAPI channel using the cell detection algorithm with the following parameters: requested pixel size 0.5 µm, background radius 8.0 µm, sigma 1.5 µm, minimum nucleus area 10 µm2, maximum nucleus area 400 µm2, intensity threshold 10, cell expansion 5.0 µm. Detected cells were classified based on mean channel intensity thresholds applied to the EpCAM, MHCII, and CD4 channels independently. Cells were classified as EpCAM^+^MHCII^+^ when mean EpCAM and MHCII channel intensities both exceeded a threshold of 10 arbitrary units. Cells negative for EpCAM, MHCII and CD4 were classified as “other”. Nearest neighbor distances from each CD4^+^ cell to the nearest EpCAM^+^MHCII^+^ cell was calculated as centroid-to-centroid Euclidean distances in µm using a custom Groovy script, which is available upon request. CD4^+^ cell density was calculated as the number of CD4^+^ detected cells per mm2 of ROI. Per-image values were averaged within each mouse to yield one value per animal for statistical comparison. All classification and proximity analyses were performed using consistent parameters across all images.

### Statistics

*Student*’s t tests or Mann-Whitney tests (when normality requirement was not met) were used for single comparisons. ANOVA was conducted for multiple group analyses. When analyzing nonparametric multiple groups, Kruskal-Wallis tests were performed with false discovery rate adjustment. The statistical analyses were performed with the aid of Prism software package. * P<0.05; ** P<0.01; *** P<0.001; ns, not statistically significant.

## Acknowledgements

We thank the Cytometry and Imaging Shared Resource (CISR) and the Biostatistics & Bioinformatics Shared Resource (BBSR) at the University of Miami Miller School of Medicine.

The authors declare no competing financial interests.

---

**Fig. S1.**
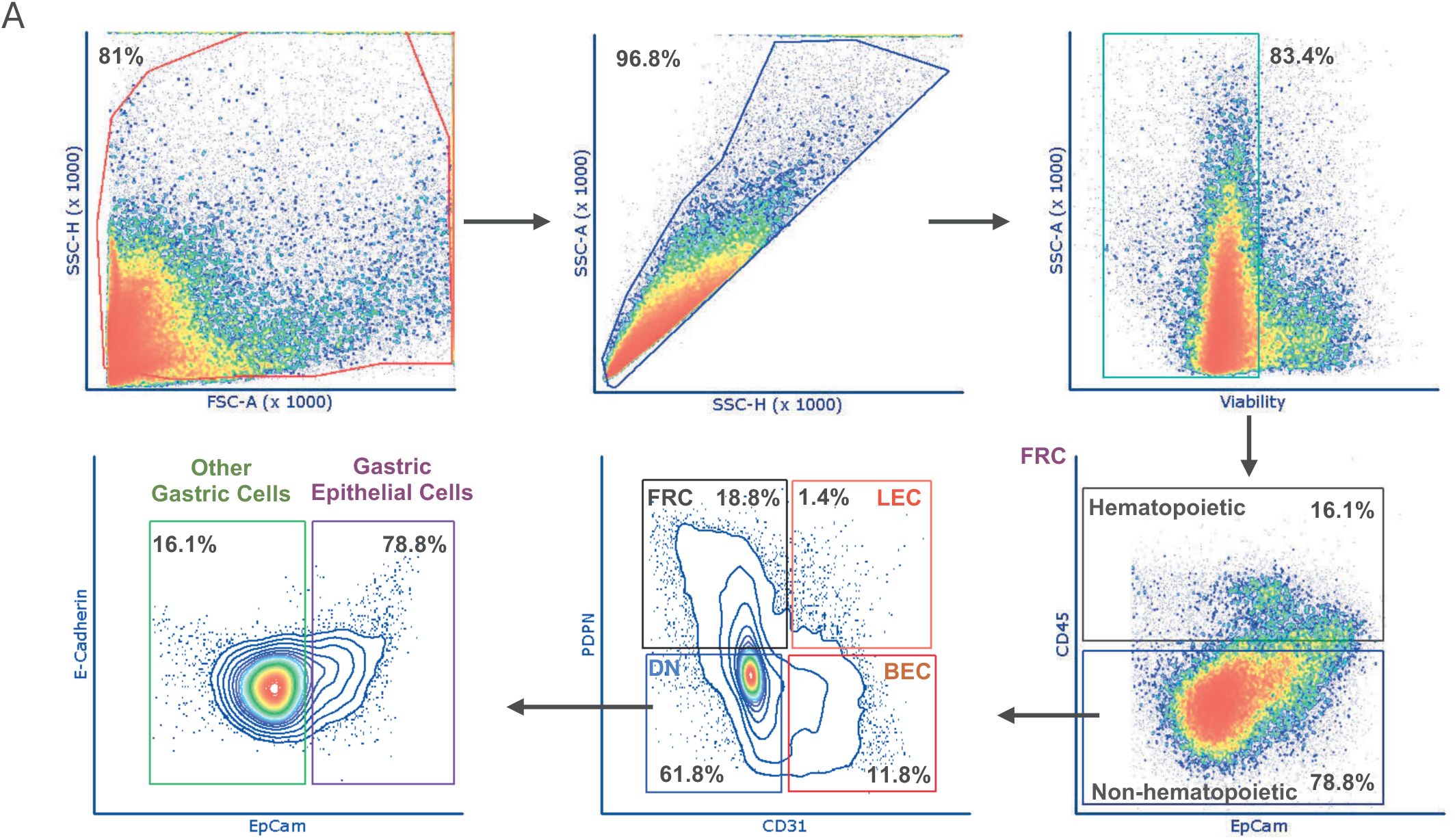
Gastric epithelial cell gating strategy by 40-color spectral flow cytometry. **(A)** Representative flow cytometry gating strategy used to identify the non-hematopoietic gastric epithelial compartment. Cells were gated on singlets, live cells, and CD45^-^. Within the CD45^-^ gate, stromal populations were defined as fibroblastic reticular cells (FRC; PDPN^+^CD31^-^), lymphatic endothelial cells (LEC; PDPN^+^CD31^+^), blood endothelial cells (BEC; PDPN^-^CD31^+^), and double-negative cells (DN; PDPN^-^CD31^-^). The gastric epithelial population was identified as DN EpCAM^+^ cells and subjected to downstream marker analysis.

**Fig. S2.**
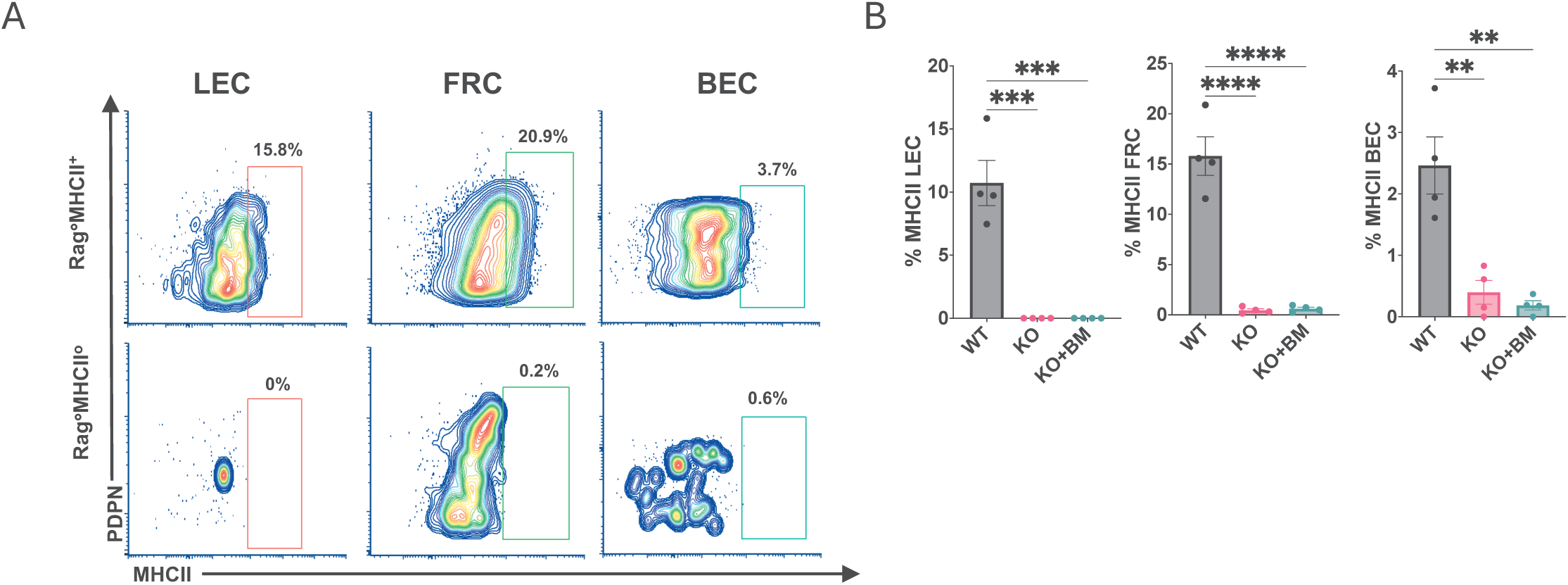
Expression of MHCII on stromal populations in the stomach. **(A)** Representative flow cytometry plots showing MHCII expression on stromal populations within the CD45^-^compartment: fibroblastic reticular cells (FRC; PDPN^+^CD31^-^), lymphatic endothelial cells (LEC; PDPN^+^CD31^+^), and blood endothelial cells (BEC; PDPN^-^CD31^+^). **(B)** Summary percentage of MHCII expression on each stromal population across experimental groups.

**Fig. S3.**
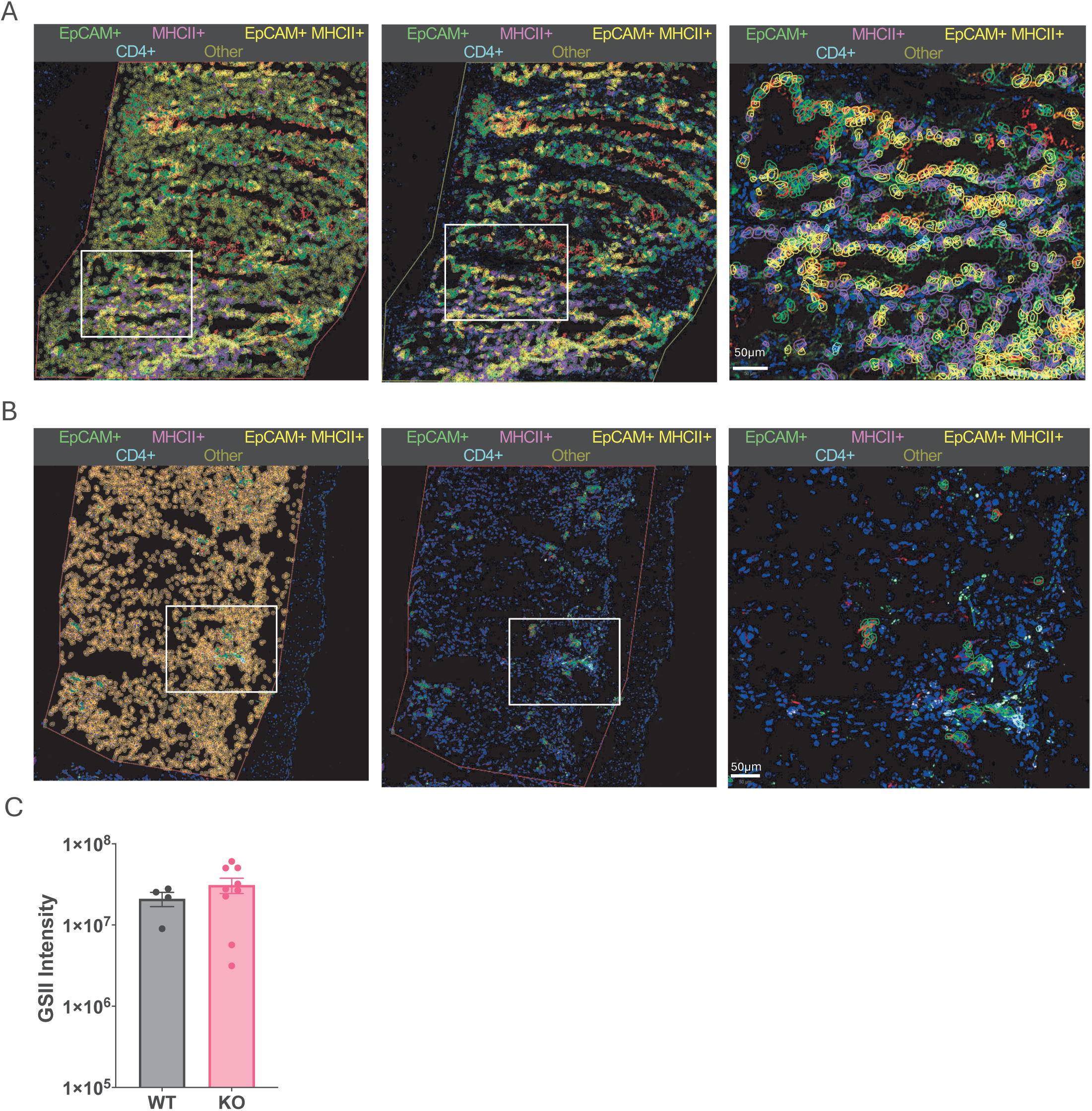
QuPath automated cell classification of gastric tissue from MHCII-wildtype and MHCII-deficient hosts. **(A)** Representative immunofluorescence image of gastric tissue from a Rag°MHCII^+^ host with automated cell classification overlay showing all detected populations: EpCAM^+^MHCII^+^ cells (yellow), CD4^+^ cells (cyan), EpCAM^+^ cells (light green), MHCII^+^ cells (magenta), and other cells (dark green). Full tissue ROI at low magnification with all classification overlays (left); Full tissue ROI at low magnification with unclassified “other” cells excluded from the overlay; magnification of the highlighted area. **(B)** Representative immunofluorescence images of gastric tissue from a Rag°MHCII° mice confirming absence of EpCAM^+^MHCII^+^ epithelial cells. Full tissue ROI at low magnification with all classification overlays (left); Full tissue ROI at low magnification with unclassified “other” cells excluded from the overlay; high magnification of the highlighted area. **(C)** Fluorescence intensity quantification of GSII within defined tissue ROIs. Corrected total cell fluorescence (CTCF) was calculated as: integrated density - (ROI area x mean background fluorescence), measured from two to five fields per tissue and averaged per mouse. Each data point represents one mouse; mean ± SEM; Mann-Whitney test. *p < 0.05, **p < 0.01, ***p < 0.001.

**Fig. S4.**
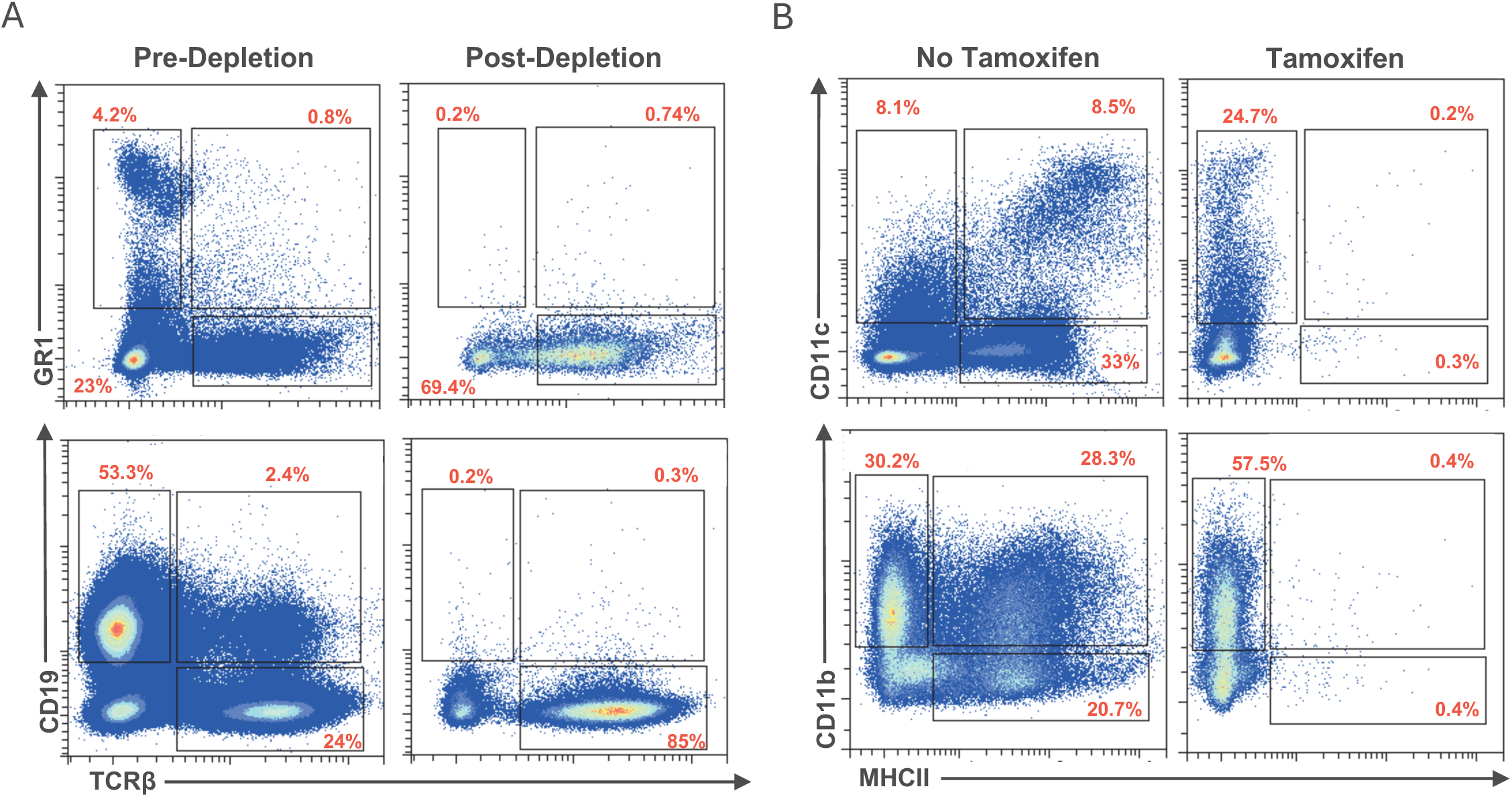
Validation of immune cell depletion and tamoxifen-induced MHCII deletion efficiency. **(A)** Representative flow cytometry analysis of depletion efficiency following magnetic bead-based removal of NK1.1^+^, CD11b^+^, CD11c^+^, F4/80^+^, Gr1^+^, and CD19^+^ cells from CTLA4KD donor splenocytes prior to adoptive transfer. Residual positive cells comprised 0.2-0.7% for GR1 and 0.2-0.3% for CD19, indicating greater than 99% depletion efficiency. **(B)** Validation of tamoxifen-induced MHCII deletion in Rag°H2Ab^fl/fl^Rosa26-CreERT2 hosts by flow cytometry analysis. Residual MHCII expression was detected in 0.2-0.4% of target cells following tamoxifen treatment, confirming 99.6-99.8% deletion efficiency.

**Fig. S5.**
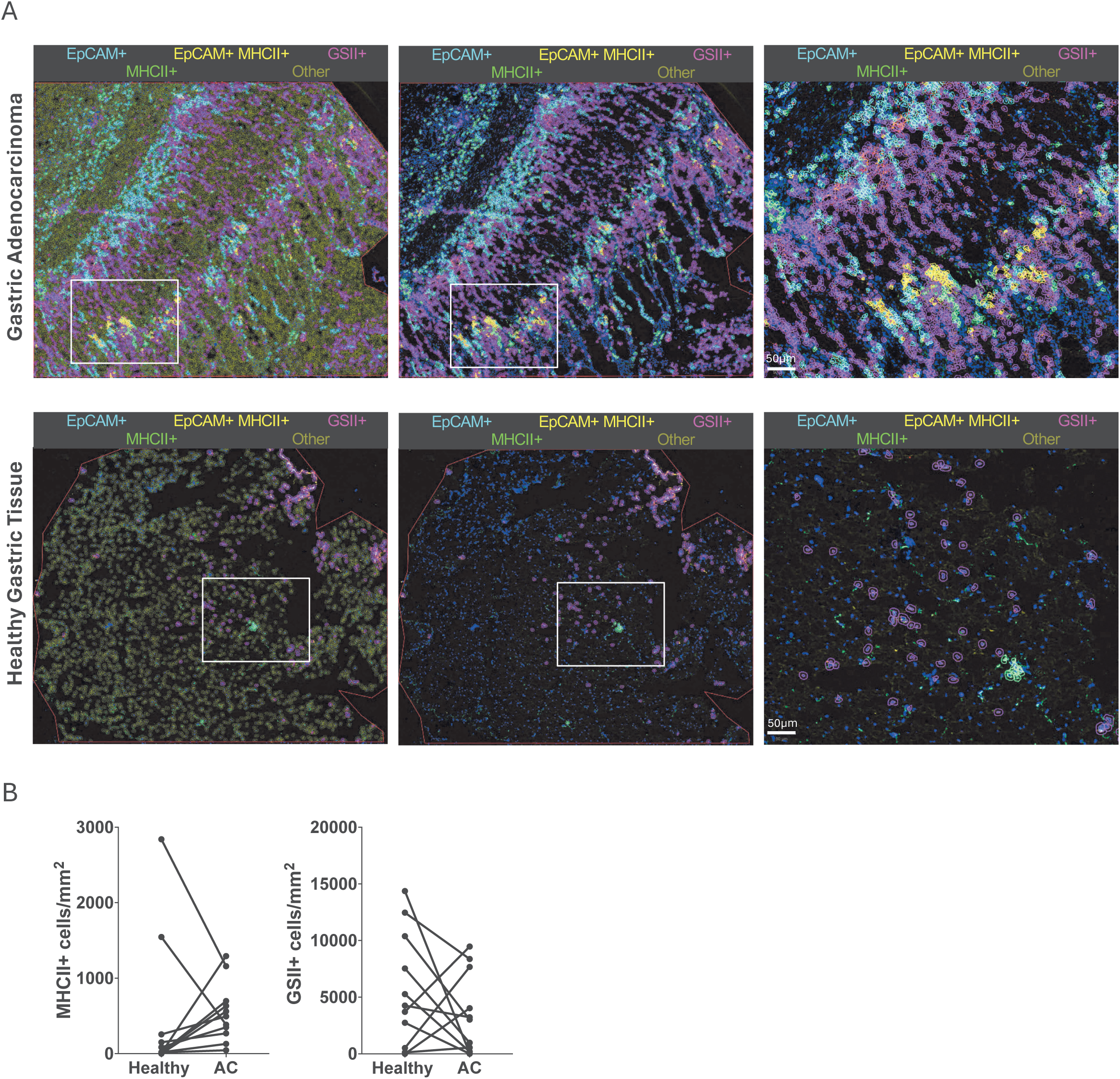
QuPath automated cell classification of human gastric adenocarcinoma and matched healthy controls. **(A)** Representative immunofluorescence image of gastric tissue from adenocarcinoma (top) and healthy controls (bottom) with automated cell classification overlay showing all detected populations: EpCAM^+^ cells (cyan), EpCAM^+^MHCII^+^ cells (yellow), GSII^+^ cells (magenta), MHCII^+^ cells (green), and other cells (dark green). Full tissue ROI at low magnification with all classification overlays (left); full tissue ROI at low magnification with unclassified “other” cells excluded from the overlay (center); high magnification of the highlighted area (right). **(B)** MHCII^+^ (left) and GSII^+^ (right) cells quantified as cells per mm2 of annotated gastric tissue ROI in human adenocarcinoma and matched healthy controls. Each point represents one patient, n=11; Wilcoxon matched pairs signed rank tests; *p < 0.05, **p < 0.01.

